# Deeper than you think: partisan-dependent brain response

**DOI:** 10.1101/2021.11.23.469735

**Authors:** Noa Katabi, Hadas Simon, Sharon Yakim, Inbal Ravreby, Yaara Yeshurun

## Abstract

Recent political polarization has highlighted the extent to which individuals with opposing views experience ongoing events in markedly different ways. In this study, we explored the neural mechanisms underpinning this phenomenon. We conducted functional magnetic resonance image (fMRI) scanning right- and left-wing participants watching political videos just before the 2019 elections in Israel. Behavioral results demonstrated significant differences between left- and right-wing participants in their interpretation of the videos’ content. Neuroimaging results revealed partisanship-dependent differences in both high-order regions and early-motor and somato-sensory regions, although no such differences were found with regard to neutral content. Moreover, we found that most of the political content was more potent in synchronizing participants with right-wing views, and that this synchronization was observed already in early visual and auditory cortices. These results suggest that political polarization is not limited to higher-order processes as previously thought, but rather emerges already in motor and sensory regions.

## Introduction

Today, perhaps more than ever, creating a shared understanding of the world we live in seems like an urgent yet elusive endeavor. Humans understand each other well enough to create social and technological feats of immense complexity, but not enough to agree on whether the media coverage of an election was biased, or in which way. In this study, we set out to test how political partisanship shape brain activation and synchronization between individuals with similar or opposing political views.

Political partisanship has been shown to influence one’s choices, perception, and understanding of information^1–3^. The divergence between how liberals and conservatives view the world also shapes their memory (e.g., participants reported remembering false events in support of their partisan views^4^) and their preferences (e.g., preferring policies suggested by their in-group politician, even if those policies are at odds with their ideology^5^). Moreover, it was found that liberals are more open to new experiences^6^ whereas conservatives have a greater need to reduce uncertainty, ambiguity, stress, and disgust^7^. Importantly, the findings suggested that conservatives are more likely to value conformity and to tend toward in-group consensus and shared reality, while liberals consider themselves less inclined to in-group consensus^8^.

These behavioral partisan-based differences were also borne out by evidence from neuroimaging studies. Previous studies demonstrated political-based differences in brain responses, as well as in the volume of specific regions^7^. For example, liberalism has been shown to be associated with increased anterior cingulate (ACC) volume, and conservatism with increased amygdala volume^9^. These extend previous neurocognitive findings about the high degree of conflict monitoring among liberals, which is associated with increased ACC activation^10^, and conservatives’ sensitivity to threatening situations and facial expressions, which reflects an emotional processing in areas such as the amygdala^11,12^. Investigating fundamental differences between people with opposing political views, it was shown that political orientation could be predicted based on the neural response to disgusting images, even after a single stimulus presentation^13^. A similar effect was also evident in a risk-taking task^14^.

These studies used relatively simple stimuli, such as static pictures, to reveal partisanship-dependent differences in brain responses. Recent studies used stimuli such as stories and movies to test for these differences. For example, using polarized political videos about immigration policy evidence for “neural polarization” (divergence in brain activity between liberals and conservatives) was found in the dorsal medial prefrontal cortex (dmPFC)^15^. Similarly, a functional near-infrared spectroscopy (fNIRS) study, in which participants watched two videos on abortion, found that the participants’ respective stances could be classified based on their response in the dmPFC^16^. Moreover, a recent study examined neural synchronization between individuals watching political and non-political video clips^17^. It found increased neural synchrony between participants who shared a political ideology during an intense pre-election political debate, while no such increased synchronization was evident during a political news video clip or, a non-political one.

In the present study, we tested how political partisanship shapes the brain response to polarizing political stimuli in a period when political partisanship was very present in everyday life, namely, just before the 2019 elections in Israel. By using functional magnetic resonance imaging (fMRI), we were able to examine neural activation and synchronization of two groups with opposing political views while they watched various political video clips. Based on previous studie,^15,18,19^, we hypothesized that there would be partisan-dependent differences in the default mode network (DMN) while processing political content. Moreover, based on behavioral studies that found conservatives have a greater tendency toward conformity^8^, we hypothesized that there would be greater neural synchronization between right-wing individuals.

## Results

In the three weeks before the April 2019 elections in Israel, 17 right-wing subjects and 17 left-wing subjects watched five videos: a neutral clip (a short documentary about a guy who moved an old bus into his house); 2 campaign ads (one by a right-wing party, and one by a left-wing party), and 2 political speeches (one by Benjamin Netanyahu, a right-wing politician, and one by Shelly Yachimovich, a left-wing politician; Fig. 1a). Following each clip, participants were asked to answer three questions about the clip they had just seen: (1) “How much did you agree with the main message of this clip?” (2) “How much did this clip interest you?” (3) “How emotionally engaged were you?”.

**Fig. 1.**
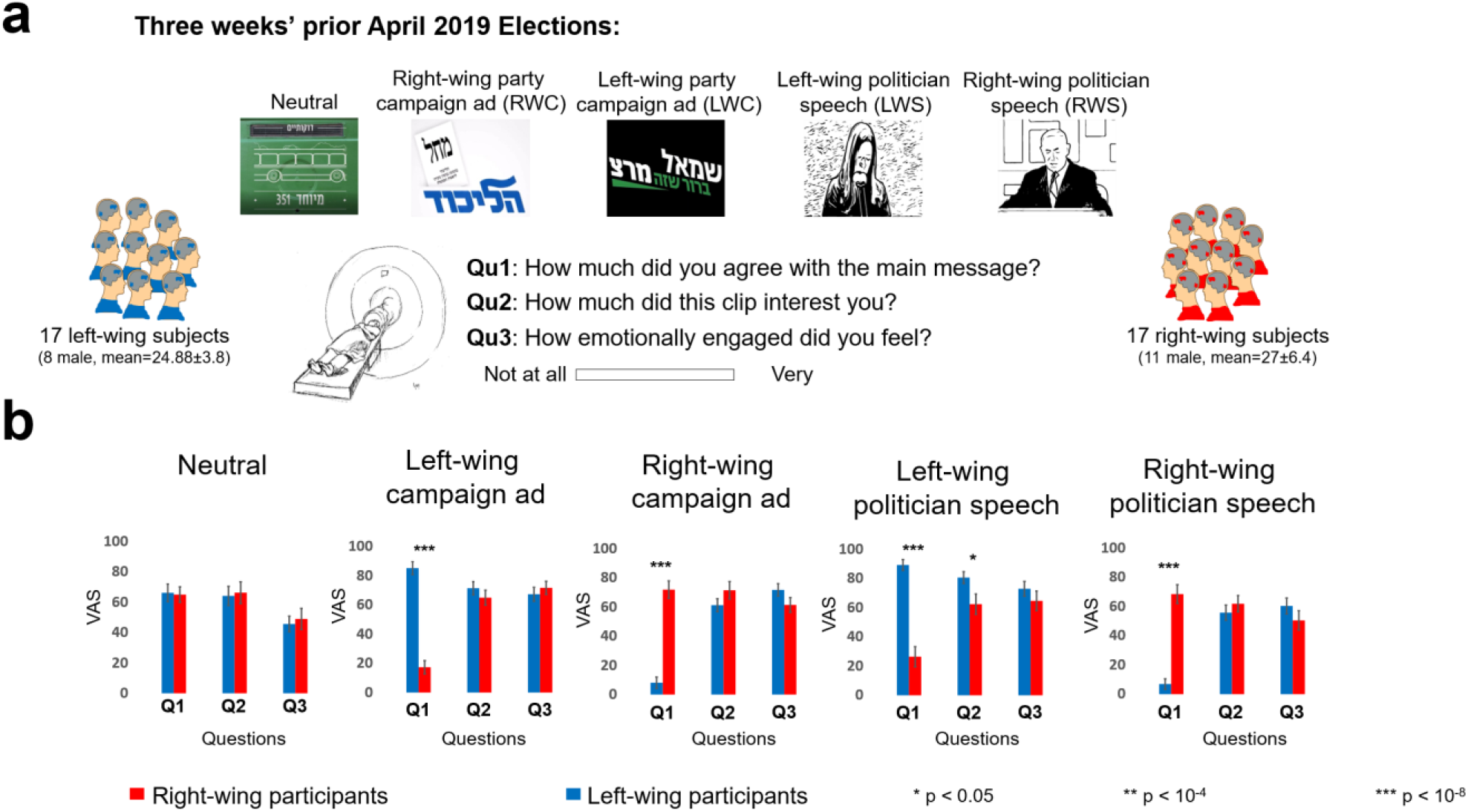
Experiment design and behavioral results. **a**, Stimuli and experiment design. Participants watched 5 video clips inside the scanner (upper panel) and answered 3 questions after each clip (lower panel). **b**, Participants’ ratings demonstrated large differences between the two groups in terms of how much they agreed with the main message of the political clips, and relatively similar emotional engagement and interest with the clips. (*p < 0.05, **\*\*** p < 10^-4^, ***p < 10^-8^; graphs—mean±ste).

### Partisanship-dependent differences in agreeing with the main messages of the video-clip

The ratings from the questions inside the scanner revealed that while the groups did not differ in the degree of their agreement with the main message of the neutral movie (L-group *M* = 66.1 *SD* = 23.8, R-group *M* = 64.97 *SD* = 20.86; t(32) = 2.03; *p* = 0.88), there was a large and significant difference in their degree of agreement with the main message of all the political video clips: right-wing campaign ad (RWC; L-group *M* = 8.14 *SD* = 15.91, R-group *M* = 71.83 *SD* = 24.77; t(32) = 2.03; *p* < 10^-9^); left-wing campaign ad (LWC; L-group *M* = 85.17 *SD* = 18.07, R-group *M* = 17.32 *SD* = 18.60; t(32) = 2.03; *p* < 10^-11^); right-wing politician speech (RWS; L-group *M* = 6.88 *SD* = 14.82, R-group *M* = 68.49 *SD* = 26.29; t(32) = 2.03; *p* < 10^-8^) and left-wing politician speech (LWS; L-group *M* = 89.19 *SD* = 15.21, R-group *M* = 26.40 *SD* = 28.28; t(32) = 2.03; *p* < 10^-8^) (Fig. 1b). Moreover, although right-wing participants were more in agreement with the main message of the RWC and RWS, and left-wing participants agreed more with the main message of the LWC and LWS, there were no significant differences in the degree to which they were interested or emotionally engaged with the clips, except with regard to the LWS, which left-wing participants found to be more interesting (L-group *M* = 80.54 *SD* = 16.49, R-group *M* = 62.35 *SD* = 28.58; t(32) = 2.03 *p* < 0.05) (Fig. 1b).

### Differentiation in regions involved in processing political content in right- and left-wing participants

We were interested in testing for partisanship-dependent brain response similarities and differences in processing the stimuli. To this end, we first divided the brain into 268 nodes, using a parcellation map based on resting state connectivity^20^. We then tested the brain response to the neutral and political clips by using inter-subject correlation (ISC) analysis^21^. Each of the five clips generated an extensive brain response among both groups, including in the sensory regions, the mentalizing network, and lateral prefrontal cortex (Fig. 2, marked in yellow).

**Fig. 2.**
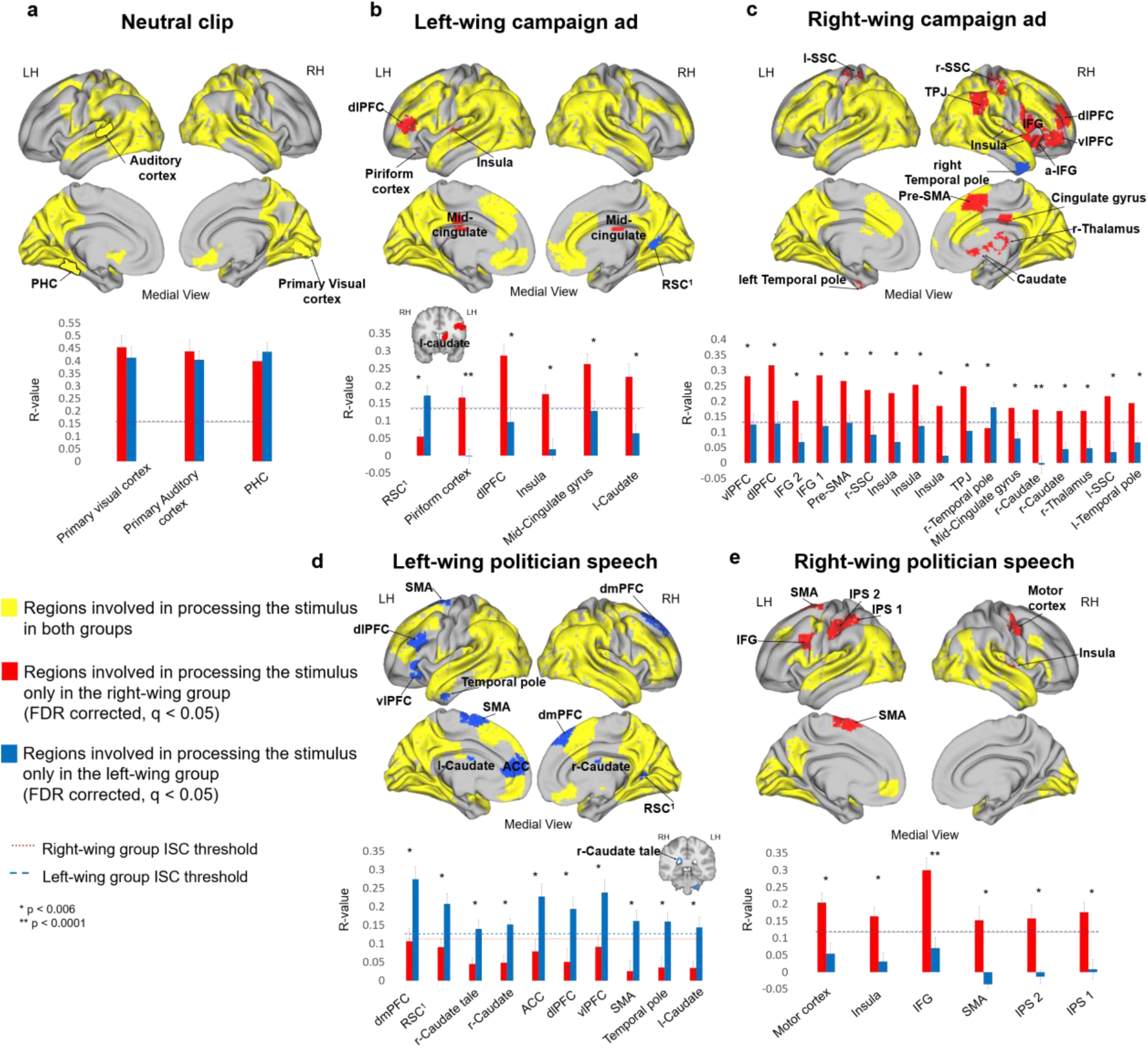
ISC results. The brain maps demonstrate regions involved in processing the stimulus in both groups (marked in yellow), in only the right-wing group (marked in red), or in only the left-wing group (marked in blue)—with regard to the neutral video clip (**a, upper panel**); the left-wing (**b, upper panel**) and right-wing (**c, upper panel**) campaigns ads; and left-wing (**d, upper panel**) and right-wing (**e, upper panel**) politician speeches (FDR corrected, q < 0.05). The graphs in the lower panel show ISC R-values of both groups in each of the “red” and “blue” brain regions (mean±ste). The dashed lines represent the group’s ISC threshold. The regions included IPS, intraparietal sulcus; IFG, inferior frontal gyrus; dlPFC, dorsal lateral prefrontal cortex; SSC, somatosensory cortex; TPJ, temporal parietal junction; vlPFC, ventrolateral prefrontal cortex; SMA, supplementary motor area; dmPFC, dorsal medial prefrontal cortex; ACC, anterior cingulate cortex; RSC, retrosplineal cortex^1^ adjacent to the RSC.

To test for the differences between the groups in terms of the brain areas involved in processing the stimulus, we performed a 2-step analysis, which yielded differences in two types of regions: regions that were involved in processing the stimuli only in one group and not in the other, and regions that were involved in processing the stimuli in both groups, but where the responses were more synchronized in one of the groups (see Methods section for details). Interestingly, we found political-group-dependent differences only with regard to the political clips, not for the neutral one (Fig. 2a). Specifically, in the case of the RWC, LWC, and RWS, we found many regions that were involved in processing the stimuli in the right-wing participants, but not in the left-wing ones (Fig. 2b,c,e, marked in red, p < 0.006; see supplementary, Table 1). These regions included the left caudate; the right- and left-dorsal lateral pre-frontal cortex (r-dlPFC; l-dlPFC); the temporal-parietal junction (TPJ) and the posterior cingulate (PCC); part of the somatosensory cortex, motor, and pre-motor areas. Only two regions were involved in processing these clips in left-wing participants, but not right-wing ones: the right temporal pole in RWC, and an area adjacent to the retrosplineal cortex (RSC) in LWC (Figure 2b-c, marked in blue, p < 0.006).

Intriguingly, this analysis revealed the opposite pattern for the LWS stimulus. Here, we found regions that were involved in processing the stimuli only within left-wing participants, but not within their right-wing counterparts (Fig. 2d, marked in blue, p < 0.006; see supplementary, table. 1). These included the dmPFC, ACC, ventrolateral prefrontal cortex (vlPFC), and the cerebellum (see supplementary, Fig. 2).

Finally, we found regions that were involved in processing the stimuli in both groups, but revealed significantly more synchronized responses within the right-wing group while watching the political campaign ads (see supplementary, table. 1). These regions included the visual cortex, the parahippocampus, and part of the intraparietal sulcus (see supplementary, Fig. 3). Taken together, these results suggest that in addition to the expected DMN and high-order regions, partisanship-dependent differences were identified already in motor and somatosensory brain regions.

### Political-view dependent neural synchrony

To further understand the political-dependent differences in neural synchronization, we applied IS-RSA^22^ to all participant responses (N = 34 in RWC and LWC; N = 32 in RWS and LWS). Specifically, we tested whether the stronger any two participants’ political views were (to the right or left), the more similar were their brain responses. For each video-clip, we generated political views similarity matrix by averaging each pair of participant’s degree of agreement with the main message (Fig. 3a), and a brain similarity matrix, by calculating the pairwise correlation between every two subjects. We then computed Spearman’s R between the behavioral matrix and the brain-similarity matrix. Similarity in brain activation was found for most of the stimuli among participants with right-wing views, particularly with regard to the political campaign ads (RWC and LWC; Fig. 3b). These regions included areas within the DMN (e.g., TPJ and PCC), dlPFC and somatosensory cortices (similar regions to those identified in the ISC results), as well as primary visual area, primary auditory cortex, insula, fusiform, sub-cortical regions (e.g., thalamus, caudate, and nucleus accumbens).

**Fig. 3.**
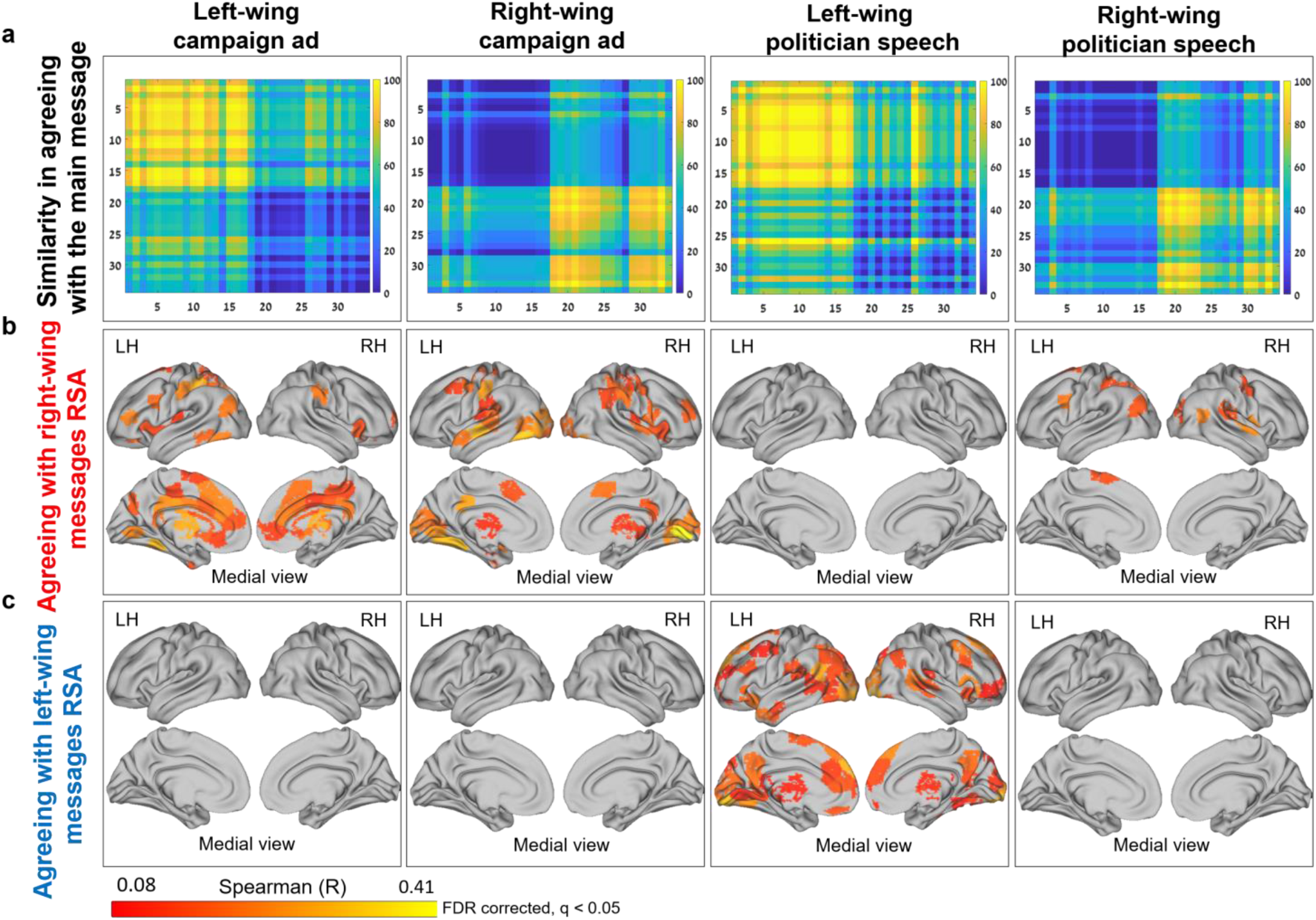
IS-RSA results: **a**) Behavioral similarity matrices based on “agreement with the main message” question; **b**) IS-RSA brain maps for agreeing with a right-wing view; **c**) IS-RSA brain maps for agreeing with a left-wing view. Red to yellow brain areas are areas with an R-value ≤ 0.407 and FDR-corrected, q < 0.05.

Moreover, in line with our ISC results, with regard to the left-wing politician speech, we found synchronized brain activation in many brain regions between people with left-wing views (Fig. 3c). These regions included the visual cortex, precuneus, dmPFC, ACC, orbitofrontal cortex and thalamus. Taken together, these results reveal that the clips’ content shaped synchronization patterns of regions within the DMN (along partisan lines), and – more surprisingly – the synchronization of early sensory and somatosensory regions as well.

### Differences between the political stimuli

To better interpret the various patterns of partisan-dependent brain-response differences with regard to the left-wing speech, we recruited 32 independent raters (16 right-wing participants, and 16 left-wing ones) to evaluate and analyze the content and goals of the four political video clips, and the participants’ reactions toward them. We analyzed participants’ responses in each group as well as across groups, and found significant differences between responses to the left-wing speech versus the other political stimuli: LWS was different from the other videos in general (53% of all raters voted for LWS; Fig. 4a) and specifically regarding its main goal - while all other stimuli focused on encouraging people to vote, LWC focused on delivering a message (97% of all raters voted for LWS; Fig. 4b). Among the left-wing raters, LWS was rated as a better representation of left-wing opinions in Israel (69% of the participants voted for LWS; Fig. 4c), and elicited less objection (all left-wing raters voted no objection; Fig. 4d) and criticism (t(15) = 4.32, p < 0.0006; Fig. 4e) than LWS.

**Fig. 4.**
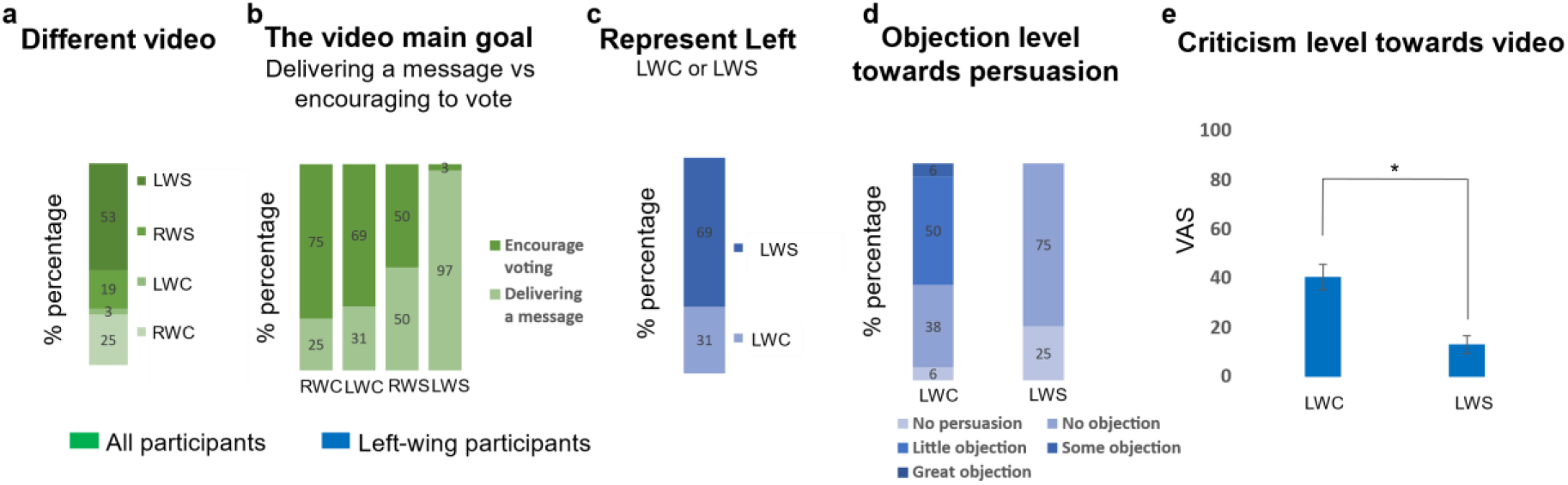
Between stimuli differences. Post-hoc experiment of 32 independent raters revealed that: **a**) the left-wing politician speech was different from the other clips—specifically in **b**) the video’s main goal; **c**, the degree to which left-wing individuals felt that it represents the left-wing perspective in Israel; and **d**, how much it evoked objection and **e**, criticism in them. p < 10^-3^.

## Discussion

Our results demonstrate partisanship-dependent differences in brain activation and synchronization of individuals processing political content. We found regions within the DMN, prefrontal areas, insula, somatosensory and motor regions that were involved in processing the political content, but only in one group, not in the other. Moreover, we found that specific political content was more potent in synchronizing the brain responses of individuals on one side of the political spectrum, but not the other. Intriguingly, these regions included early sensory regions, such as V1 and A1. Significantly, these activation and synchronization differences occurred only with regard to political content, and not found when the participants were processing a neutral stimulus.

Our results revealed that certain regions within the DMN – such as the TPJ, PCC and dmPFC – were uniquely involved in processing the political stimuli among the right-wing group (while watching both campaign ads and the right-wing politician speech) and in the left-wing one (while watching the left-wing politician speech). This is in line with prior studies that used non-political narratives and found that the DMN was involved in comprehension and interpretation of naturalistic stimuli^18,19,23,24^. Moreover, recent studies suggested that dmPFC and TPJ responses differed between conservatives and liberals while they watched polarized political content^16,17^, and that this difference increased when emotional language was involved^15^. Alongside the DMN, the lateral prefrontal cortex, including the dlPFC, was involved in processing political campaign ads, but only in the right-wing group. This result is consistent with previous findings of the dlPFC neural response as a predictor of conservative ideology^13^. It is worth noting that the dlPFC activation in the right-wing group was observed with regard to both right-wing and left-wing campaign ads—suggesting that the involvement of the dlPFC is in line with the role of the prefrontal cortex in emotion regulation and appraisal when the subject is expecting to be asked about their views on certain stimuli or their interpretation of them^25^.

Interestingly, our results revealed that the somatosensory, pre-motor and motor regions were involved in processing both campaign ads and the right-wing politician speech only in right-wing individuals (Fig. 2b,c,e). We would like to interpret these findings through the lens of *embodied cognition*^26^, simulation theory^27^, and the bi-directional link between body movements and cognition^28^. It has been suggested that the sensorimotor experience is part of how an event is represented in the brain^29^, as demonstrated by findings that people use sensorimotor representation to process action and non-action words and sentences^30,31^ as well as facial expressions^32^. Moreover, it was suggested that such simulative representation mechanisms offer an adaptive value to understanding other people’s actions and intentions^33,34^. Our results suggest that right-wing individuals (but not left-wing ones) use sensorimotor simulative representation to process political content, which, in turn, may facilitate their identification with the right-wing content, through a mechanism similar to that found in other studies among people who felt enhanced empathy for their fellow in-group members who were experiencing pain^35,36^.

Partisanship-dependent differences also emerged in the groups’ neural synchronization. Using IS-RSA, we found that most of the political stimuli were more potent in synchronizing the brain responses of individuals with right-wing views, while the left-wing politician speech was more potent in synchronizing the brain responses of their left-wing counterparts (Fig. 3). Notably, this pattern was striking in its dichotomy: agreeing with a right-wing message presented in most of the clips synchronized the participants’ responses in many different brain regions, whereas in the same clips, agreeing with a left-wing message did not synchronize participant responses in any brain region (and vice versa for the left-wing politician speech). Specifically, the results revealed that the more two individuals share a right-wing outlook while watching the campaign ads and the right-wing speech, or a left-wing outlook while watching the left-wing politician speech, the more synchronized their neural responses were in various regions of their brain—including the DMN, dlPFC, Parahippocampus, somatosensory regions, primary visual and auditory cortices, and the subcortical areas (e.g., thalamus, caudate and, nucleus accumbens; Fig. 3). While these highly synchronized responses of the DMN and lateral pre-frontal cortex are in line with previous studies that found that these regions are involved in subjective interpretation^18,24,37–39^ – specifically in political contexts^15,17,40^ – we would like to expand upon the unexpected increase in partisanship-dependent synchronization in the insula and primary sensory cortices.

The insula was shown to be involved in affective processing and emotional engagement^41,42^, including feelings such as disgust^43^, as well as envy and resentment toward competitive members^44^. It was also found to be more activated when viewing a candidate from the opposite political party, especially when Republicans viewed a Democratic candidate^45^. Our results suggest that the campaign ads that were used in this experiment were potent in synchronizing responses in brain areas that reflect the emotional engagement participants with right-wing views experienced while watching the right- and left-wing political campaign ads, and maybe even resistance and disgust, which were elicited by the political campaign ad of the opposing side.

Surprisingly, our results revealed greater partisanship-dependent synchronization in primary visual and auditory cortices. In other words, the political views of individuals shape their neural responses at a very basic level: they are seeing and hearing different things not just metaphorically, but rather literally. This is in contrast to previous studies, that found similar between-groups activation in primary sensory regions regardless of whether participants understood the narrative or not^23,46^, or understood it in different ways^15,17,18,40^. We suggest that the timing of this experiment – just three weeks before the elections in Israel, when the political atmosphere was tense and fraught – had increased the participants’ engagement and emotional reaction to the stimuli, thereby revealing effects that had not previously been demonstrated.

Our results revealed a distinct pattern of activation and synchronization that was dependent on the specific content: three of the four political video clips (two political campaign ads, and a right-wing politician’s speech) evoked increased activation and synchronization within right-wing individuals, while the left-wing politician’s speech generated increased activation and synchronization within left-wing individuals. The former result is in line with previous findings that conservatives have a strong tendency to in-group consensus, conformity, and shared reality with like-minded people^8^, while liberals tend more toward intragroup variability and within-group differences^47,48^. To better understand what made the left-wing politician’s speech unique in inducing synchronization among left-wing individuals, we ran a behavioral experiment. We found that 97% of 32 independent right- and left-wing raters thought that the main goal of the left-wing politician’s speech was to deliver a message, while the other stimuli were simply encouragements to vote; that the left-wing political campaign ad (but not the politician speech) rankled in its delivery and content among the left-wing raters; and that the left-wing politician’s speech (but not the left-wing political campaign ad) was the clip that was deemed to be the best fit to left-wing views in Israel (Fig. 4). These attributes may explain the less pronounced left-wing neural synchronization in response to the left-wing campaign ad, and the greater synchronization to the left-wing politician’s speech. Taken together, we suggest that individuals with a right-wing outlook are more likely to synchronize with each other – both when they agree and when they don’t agree with the message – due to their higher tendency to in-group conformity, whereas left-wingers’ individualism and self-criticism possibly reduce in-group synchronization.

It is evident that the political sphere has become highly polarized in recent years^49^. Our finding of partisanship-dependent differences in activation and synchronization already in primary sensory and motor regions may contribute to our understanding of how such differences come about. We hope that these findings could generate fruitful pathways for future research. Specifically, we suggest examining whether top-down or bottom-up mechanisms generate such partisanship-dependent differences in primary sensory cortices, a theoretical insight that could support the development of new methods to attenuate processes of increased polarization.

## Methods

### Participants

Forty-one right-handed participants took part in this fMRI study (24 males and 17 females, mean age = 26.5±5.75). Prior to taking part in the study, they completed questionnaires about their political views, and only individuals who reported strong left- or right-wing political views were recruited for the experiment. Seven participants were discarded from the analysis: 4 due to our inability to characterize their political views based on their post-scan questionnaire; 2 due to incidental clinical findings; and 1 due to excessive head motion (>2mm). The remaining 34 participants were divided into two equal groups based on their political views: *Right-wing* (R group; 11 males and 6 females, mean age = 27±6.4) and *Left-wing* (L group; 8 males and 9 females, mean age = 24.88±3.8) groups. This sample size of 17 participants in each group has been shown in previous studies to be sufficient to test for similarities and differences in neural responses to naturalistic stimuli^18,23^, and for power analyses of inter-subject correlation^50^. In two (of the five) stimuli we analyzed, we excluded a further two participants due to lack of data. The experimental procedures were approved by Tel Aviv University’s Ethics Committee and the Institutional Review Board at the Sheba Tel-Hashomer Medical Center. All participants provided written informed consent, and received payment for their time.

### Stimuli and experimental design

The experiment took place three weeks before the April 2019 national elections in Israel. The scanner session involved the use of 8 video clips (mean length = 197 seconds, *SD* = 64.92 seconds): one neutral clip (a short documentary about someone who moved an old bus into his house); 4 campaign ads (2 from a right-wing, 1 from a center, and 1 left-wing party, respectively); 2 political speeches (one by Benjamin Netanyahu, a right-wing politician, and one by Shelly Yachimovich, a left-wing politician), and a pre-election political survey clip. Each clip was preceded and followed by a gray screen: 8 seconds before, and 10 seconds after (which were discarded from the analysis). After each clip, participants were asked to answer three questions about it: (1) *“How much did you agree with the main message of this clip?’’;* (2) *“How much did this clip interest you?’’;* and (3) *“How emotionally engaged were you?”*. They answered these questions by indicating their ratings (using a magnet-compatible mouse) on a visual analog scale (VAS). The order of the clips was randomized into 4 versions, which all began with the neutral clip and ended with the political survey one. The participants’ eye gaze was monitored online and recorded at 500 Hz using SR-Research’s EyeLink 1000 Plus Eye-Tracker. However, due to technical issues, eye-tracking data was not collected in over half of the participants (18 out of 34 participants), so we did not analyze that data. Immediately after the scan, participants completed a behavioral assessment session, that was held in a separate, adjacent room (see supplementary, Fig. 1).

In this study, we analyzed the brain responses to five of the eight video clips: the neutral one (Neutral, 151 seconds long); one right-wing campaign ad (RWC, 198 seconds long); one left-wing campaign ad (LWC, 206 seconds long); a right-wing politician’s speech (RWS, 235 seconds long); and a left-wing politician’s speech (LWS, 312 seconds long).

### MRI acquisition

Participants were scanned using a 3T Siemens Prisma scanner with a 64-channel head coil. T1-weighted structural images were acquired using a magnetization prepared rapid gradient echo pulse sequence (MPRAGE), as follows: TR=2530ms, TE=2.88ms, TI=1100ms, flip angle= 7° and 250Hz/px, isotropic voxel size of 1mm^3^. For Functional scans, images were acquired by means of a T2*-weighted multiband echo planar imaging protocol. Repetition time (TR) = 1000 ms, echo time (TE) = 34ms, flip angle (FA) = 60°, multiband acceleration factor of six without parallel imaging. Isotropic resolution was 2mm^3^ (no gaps) with full brain coverage; slice-acquisition order was interleaved.

### Imaging analysis

#### Preprocessing

Raw DICOM format imaging data was converted to NIfTI with dcm2nii tool. The NIfTI files were organized according to the BIDS format v1.0.1^51^. fMRI data preprocessing was conducted using the FMRIB’s Software Library’s (FSL v6.0.2) fMRI Expert Analysis Tool (FEAT v6.00)^52^. All data was subjected to the following preprocessing procedures: brain-extraction for skull-stripping anatomy image; slice-time correction; high-pass filtering (2 cycles per stimulus’s length); motion-correction to the middle time-point of each run; and smoothing with a 4-mm FWHM kernel. All images were registered to the high-resolution anatomical data using boundary-based reconstruction, and normalized to the Montreal Neurological Institute (MNI) template, using nonlinear registration. Blood-oxygen-level-dependent (BOLD) response was normalized (z-scored) within subjects for every voxel for each video-clip. Hemodynamic response function (HRF) was calculated for each participant according to the peak start time of the BOLD response in early auditory areas (A1+), using the pre-election political survey video clip (which was not part of the stimuli analyzed in this paper). The shift was then calculated as the duration from the stimulus onset to the first peak of the hemodynamic response in A1+ (mean shift = 3.18 s, *SD* = .72). We further analyzed the data only in voxels that had a reliable BOLD signal (<3000 AU) in at least 90% of the participants in each group (right-wing and left-wing). This procedure resulted in 214,919 voxels for the RWC video clip; 217,149 voxels for the LWC video clip; 217,232 voxels for the RWS; 217,305 voxels for the LWS; and 217,930 voxels for the neutral video clip. Next, we divided the brain into 268 nodes, including cortical and sub-cortical regions, using a parcellation map based on resting state connectivity^20^. Within each node, we averaged the BOLD responses of all reliable voxels of each participant, for each video-clip.

#### Inter-subject correlation

We used inter-subject correlation^21^ (ISC) to define nodes that responded reliably to each of the video clips (Neutral, RWC, LWC, RWS, and LWS). ISC measures the degree to which neural responses to naturalistic stimuli are shared between participants processing the same stimuli. For each node, we correlated each participant’s time course with the average time-course response across all other participants in the same group. We then averaged these 17 correlations values (or 16 values for RWS and LWS) to get an estimate of the in-group similarity in neural response for each of the 268 nodes. To determine whether a specific ISC value was significantly greater than chance, we calculated a null distribution generated by a bootstrapping procedure. For each of the video clips, for every empirical time course at every node, 1,000 bootstrap-time series were generated using a phase-randomization procedure. Phase randomization was performed by fast-Fourier transformation (FFT) of the signal, randomizing the phase of each Fourier component, and then inverting the Fourier transformation. This procedure leaves the power spectrum of the signal unchanged, but removes temporal alignments of the signals. Using these bootstrap-time courses, a null distribution of the average correlations was calculated for each node.

To correct for multiple comparisons, we selected the highest ISC-value from the null distribution in each node^53^. The chosen threshold for every group, in each stimulus, was defined as the top 5% of the maximum values in the 268 nodes (RWC threshold ISC: R group > 0.134, L group > 0.137; LWC threshold ISC: R group > 0.142, L group > .141; RWS threshold: R group > 0.125, L group > 0.127; LWS threshold: R group > 0.115, L group > 0.129; Neutral threshold ISC: R group > 0.151, L group > 0.154). These thresholds were used to test for regions that were involved in processing the video clips in each partisan group.

#### Testing for between-groups differences in processing the stimuli

To test for differences between the groups in neural processing of the video clips, for each of the 5 video clips we tested for nodes in which either or both of the groups had passed their ISC threshold. Next, we calculated the ISC value of each participant in each group (i.e., correlation between participant’s response to the mean response of the group). We then applied Fisher transformation to these correlation coefficients and computed a two-sample t-test between the groups (i.e., comparing 17 values of right-wing participants to 17 values of left-wing participants). This resulted in a p-value for each node. To correct for multiple comparisons, false discovery rate^54^ (FDR) correction was applied with q criterion < .05. This analysis identified two types of regions: (i) regions that were involved in processing the stimuli only in one group and not in the other, and (ii) regions that were involved in processing the stimuli in both groups, but were significantly more synchronized in one of the groups.

#### Inter-subject representational similarity analysis (IS-RSA)

We used IS-RSA^22^ to test whether similarity of political views was reflected in the similarity of brain responses to the political video clips. The similarity of political views was calculated using the AnnaK model^22^. For each video clip, and for each pair of participants, we calculated their averaged agreement value with the main message of the video clip during the scan (VAS, ranging from *0* = Disagree to *100* = Agree). This resulted in a 34×34 similarity matrix for the RWC and LWC, and a 32×32 similarity matrix for the RWS and LWS. For the brain activity similarity matrix, we used the ISC method and computed a pairwise Pearson correlation between each pair of participants’ time courses, for each node (for RWC and LWC: 34×34 brain activity similarity matrices; for RWS and LWS: 32×32 brain activity similarity matrices). The IS-RSA was then computed as the Spearman correlation between the behavioral matrix and the brain activity similarity matrix.

To determine whether a specific IS-RSA value was significantly greater than chance, we calculated a null distribution generated by a bootstrapping procedure. For each video clip, in each node, we generated a pseudo neural similarity matrix by means of a phase-randomization procedure (for each pair, correlating one participant’s randomized timecourse with the other participant’s intact timecourse – as with the ISC null distribution). We then computed the Spearman’s correlation between the “pseudo” similarity brain matrix and the behavioral matrix. We repeated the phase randomization procedure 1,000 times, to generate a null distribution of pseudo IS-RSA values for each node. The p-values of the empirical Spearman’s R-values were computed using the following formula: (number of null values larger than the real value + 1)/1,000. To correct for multiple comparisons, FDR correction^54^ was applied, with q criterion < .05.

#### Post-hoc analysis of stimuli’s content

Following our neuroimaging results, we wanted to better characterize the content of the five video clips we used in this study. Accordingly, we recruited 32 independent raters: 16 with left-wing political views (8 women, 8 men, mean age =25.31±2.72) and 16 with right-wing political views (8 women, 8 men, mean age = 22.87±2.36). They watched all five video clips in 4 different orders (but always starting with the neutral clip), and answered 18 questions on each one (VAS, and forced-choice) regarding its content, and message. These included: *“To what extent do you think the video is about persuading you to vote?”; “What do you think is the main aim of the video?”; “To what extent does the video provokes criticism in you?”;* and *“To what extent do you think that the video is representative of the values of the political party it belongs to?”* In addition, we posed three general questions about all four political video-clips: *“Of all the political video clips, which of them is different from the others?”; “Which of the two left-wing video clips best represents the left-wing public in Israel?”*; *“Which of the two right-wing video clips best represents the right-wing public in Israel?”*

## Acknowledgments

We would like to thank Yohay Zvi, Dvir Caspi and Muhammad Badarnee for their assistance in scanning and data analysis. This research was supported by the Israel Science Foundation (grant No. 2434/19).

