## Supplemental Materials for "Deeper than you think: partisan-dependent brain response"

**Supplementary materials**

Behavioral assessment

Following the scanner session, all subjects completed 9 questionnaires, one for each video-clip they saw and one about their general feelings and opinions about the politicians presented in the video-clips. The video-clips questionnaires were presented on a computer screen and ordered as presented in the scanner. They had in total 90 visual analog-scaled (VAS) ratings and forced-choice questions on both comprehension (‘political-view independent’) and interpretation (‘political-view dependent’) of the clips. Out of all video-clips questionnaires, 61 questions were political-view dependent (e.g., "To what extent do you believe that the legal proceeding will prove that Benjamin Netanyahu is innocent?") and 29 were political-view independent (e.g., "Is it argued in the video that 'unemployment is at an all-time low'?").

Behavioral results

The post-scan questionnaires revealed no significant differences between the groups in comprehension (L-group: *M* = 88.23, *SD* = 8.31, R-group: *M* = 85.29, *SD* = 14; t(12) = 2.44, p = 0.73)(Fig, 1B) or interpretation (L-group: *M* = 26.94, *SD* = 9.74, R-group: *M* = 24.71, *SD* = 12.3, t(12) = 2.17, p = 0.7) of the neutral video-clip (Fig. 1A).

In line with our expectations, the 2 partisan groups significantly differ in how they interpret the political video clips: RWC: (L-group *M* = 19.64, *SD* = 18.48, R-group *M* = 56.16, *SD* = 12.83; t(14) = 2.14, p = 0.0004), LWC: (L-group *M* = 61.99, *SD* = 16.99, R-group *M* = 38.98, *SD* = 16.15; t(12) = 2.17, p = 0.02), RWS: (L-group *M* = 31.25, *SD* = 17.43, R-group *M* = 61.53, *SD* = 2.01; t(4) = 2.77, p = 0.04) and LWS: (L-group *M* = 69.29, *SD* = 13.45, R-group *M* = 33.59, *SD* = 16.06; t(6) = 2.44, p = 0.01) (Fig. 1A). This was true for all interpretation questions, except one question in the right-wing politician speech – “in your opinion, when Benjamin Netanyahu talked about the suffering his family had been through in the last three years, did he really was excited?” (L-group *M* = 51.3 *SD* = 29.5, R-group *M* = 62.06 *SD* = 30.92); t(30) = 2.04; *p* = 0.32).

Moreover, these questionnaires revealed that the groups did not significantly differ in their performance in the political-view independent questions (Fig. 1B).

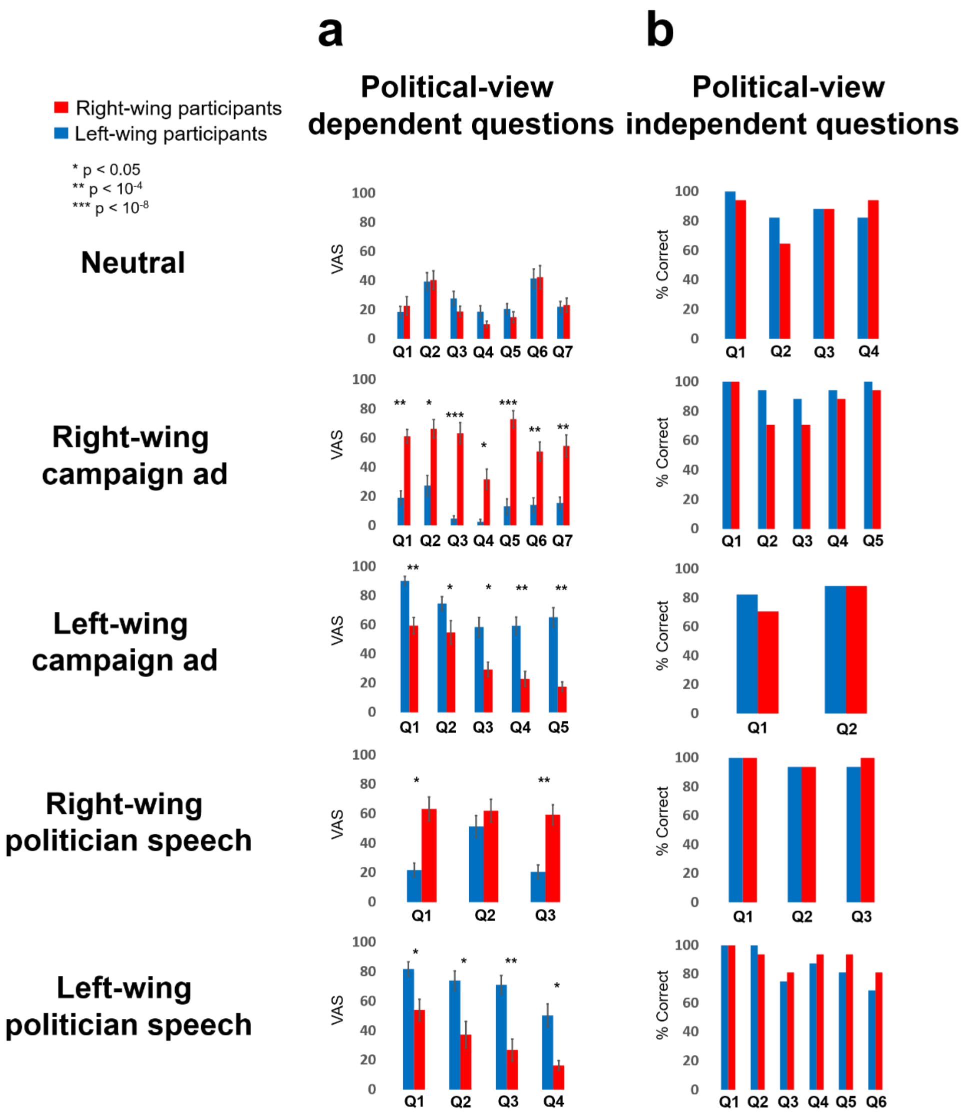


**Fig. 1.** Behavioral results demonstrating the political-view effect. **a**, The political-view dependent questions from the questionnaires of each stimulus. Significant differences were found in every interpretation question only in the political stimuli, except of one question in RWS. **b**, The political-view independent questions, as we expected, did not differ between the political groups. (*$p < 0.05$, ** $p < 10^{-4}$, ***$p < 10^{-8}$; graphs – mean±ste).

### ISC results - Cerebellum

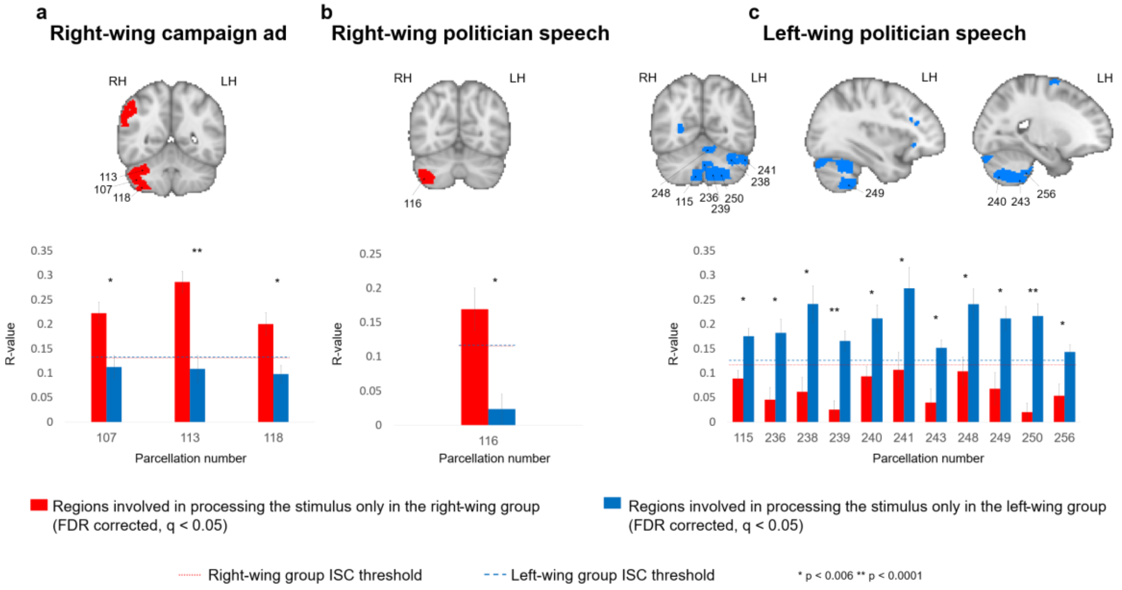


**Fig. 2.** ISC results – Cerebellum. All parcellations (Shen et al., 2013) of the Cerebellum that were involved only in one of the groups and was significantly more correlated (red for the right-wing group, blue for the left-wing group, FDR corrected, q < 0.05). The graphs indicate the R-values of the groups in every significant node (bars – mean±ste). The dashed lines are the group's ISC threshold.

### ISC results – regions that were involved in processing the stimuli in both groups but had significantly more synchronized responses in one of the groups

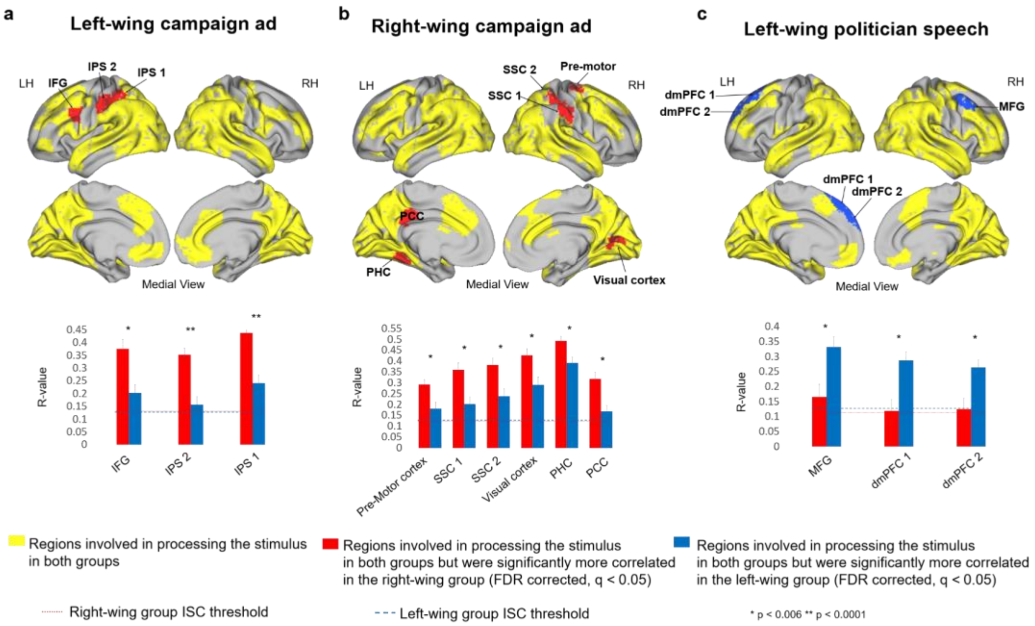


**Fig. 3.** ISC results. The red and blue brain areas are those who were involved in processing the stimulus in both groups and were significant more correlated in the R group or L group (respectively; FDR corrected, q < 0.05). The graphs indicate the R-values of the groups in every significant node (bars – mean±ste). The dashed lines are the group's ISC threshold. IFG, inferior frontal gyrus; IPS, intraparietal sulcus; SSC, somatosensory cortex; PHC, parahippocampus; PCC, posterior cingulate cortex; dmPFC, dorsal medial prefrontal cortex; MFG, medial frontal gyrus;

| | Brain region | Parcel number | Right-wing group R-value | Left-wing group R-value | p value |
|---|---|---|---|---|---|
| Right-wing campaign ad | ventro-lateral Prefrontal cortex (vlPFC) | 8 | 0.281663618 | 0.124647609 | 0.005740883 |
| | dorsal lateral Prefrontal cortex (dlPFC) | 9 | 0.317106272 | 0.127869051 | 0.000680215 |
| | inferior Frontal gyrus (IFG) 2 | 16 | 0.201838607 | 0.067854314 | 0.000997031 |
| | inferior Frontal gyrus (IFG) 1 | 21 | 0.284261697 | 0.11984986 | 0.000177213 |
| | Pre-Supplementry Motor cortex (pre-SMA) | 28 | 0.265853675 | 0.129722755 | 0.001480891 |
| | right-Somatosensory cortex | 33 | 0.236741798 | 0.091950018 | 0.000249812 |
| | Insula | 34 | 0.226793931 | 0.06793036 | 0.001001179 |
| | Insula | 36 | 0.253632625 | 0.120036147 | 0.004058332 |
| | Insula | 37 | 0.185117948 | 0.023285424 | 0.000219385 |
| | Temporal parietal junction | 47 | 0.248938271 | 0.104259211 | 0.001838113 |
| | right Temporal pole | 57 | 0.112859953 | 0.180711552 | 0.005745508 |
| | mid-Cingulate gyrus | 88 | 0.178596062 | 0.079448883 | 0.001986054 |
| | right-Caudate nucleus | 121 | 0.173126603 | -0.005745837 | 2.03E-05 |
| | right-Caudate nucleus | 123 | 0.168722399 | 0.045106209 | 0.000323011 |
| | right Thalamus | 128 | 0.168996137 | 0.048147906 | 0.002678115 |
| | left-Somatosensory cortex | 167 | 0.216761205 | 0.035082372 | 0.000567027 |
| | left Temporal pole | 202 | 0.194293245 | 0.066491606 | 0.001402024 |
| | Cerebellum | 107 | 0.222779931 | 0.112676192 | 0.002041348 |
| | Cerebellum | 113 | 0.286654288 | 0.10891178 | 1.92E-05 |
| | Cerebellum | 118 | 0.200593547 | 0.098199341 | 0.001361587 |
| | **Pre-Motor cortex** | **26** | **0.292530619** | **0.181054906** | **0.005665347** |
| | **Somatosensory cortex 1** | **38** | **0.360109424** | **0.202224634** | **0.001793549** |
| | **Somatosensory cortex 2** | **45** | **0.381966359** | **0.238585623** | **4.26E-03** |
| | **Visual cortex** | **82** | **0.426501239** | **0.290809337** | **0.00563985** |
| | **Parahippocampus gyrus (PHC)** | **198** | **0.492957399** | **0.391656067** | **0.004953297** |
| | **posterior Cingulate cortex (PCC)** | **223** | **0.318108321** | **0.16844128** | **0.000839135** |
| Left-wing campaign ad | adjacent to the Retrosplineal cortex (RSC) | 87 | 0.054760134 | 0.172599501 | 0.001834386 |
| | Piriform cortex | 135 | 0.166635939 | -0.00164614 | 8.84E-05 |
| | dorsal lateral Prefrontal cortex (dlPFC) | 154 | 0.286743133 | 0.09685222 | 1.35E-03 |
| | Insula | 170 | 0.176572929 | 0.018787769 | 0.000530824 |
| | Mid-Cingulate cortex | 224 | 0.26267003 | 0.128300996 | 0.001866731 |
| | l-Caudate nucleus | 258 | 0.2261971 | 0.064225399 | 0.000982825 |
| | **inferior Frontal gyrus (IFG)** | **157** | **0.376030084** | **0.203038404** | **1.33E-03** |
| | **Intraparietal sulcus (IPS) 2** | **171** | **0.352879518** | **0.156949755** | **2.74E-05** |
| | **Intraparietal sulcus (IPS) 1** | **179** | **0.438333498** | **0.241452459** | **5.03E-05** |
| Right-wing politician speech | Motor cortex | 27 | 0.204692104 | 0.054304817 | 0.001848954 |
| | Insula | 37 | 0.164546285 | 0.031492065 | 0.001528941 |
| | inferior Frontal gyrus | 157 | 0.300264174 | 0.071035343 | 3.81232E-05 |
| | Supplementary Motor cortex (SMA) | 162 | 0.152740245 | -0.035782314 | 0.000230684 |
| | Intraparietal sulcus (IPS) 2 | 171 | 0.15823344 | -0.013339452 | 0.000640032 |
| | Intraparietal sulcus (IPS) 1 | 179 | 0.176638418 | 0.008657955 | 0.000235378 |
| Left-wing politician speech | right dorsal medial Prefrontal cortex (dmPFC) | 12 | 0.106593352 | 0.27403593 | 0.001365341 |
| | adjacent to the Retrosplineal cortex (RSC) | 87 | 0.090844607 | 0.20754255 | 0.002105866 |
| | Caudate nucleus tale | 120 | 0.045221768 | 0.139963159 | 0.003109876 |
| | right-Caudate nucleus | 122 | 0.048601543 | 0.151721747 | 0.000310526 |
| | anterior Cingulate cortex (ACC) | 140 | 0.079249108 | 0.227382029 | 0.004050548 |
| | dorsal lateral Prefrontal cortex (dlPFC) | 147 | 0.05103565 | 0.193957963 | 0.005359024 |
| | ventro-lateral Prefrontal cortex (vlPFC) | 151 | 0.09153541 | 0.238192891 | 0.003260951 |
| | Supplementary Motor cortex (SMA) | 162 | 0.025724141 | 0.16133073 | 0.002964581 |
| | Temporal pole | 194 | 0.035776768 | 0.159820123 | 0.002889317 |
| | left Caudate nucleus | 260 | 0.034555397 | 0.143986683 | 0.003256899 |
| | Cerebellum | 115 | 0.089186464 | 0.175682416 | 0.000766296 |
| | Cerebellum | 236 | 0.045861221 | 0.182617233 | 0.000914433 |
| | Cerebellum | 238 | 0.062093042 | 0.241649461 | 0.000741513 |
| | Cerebellum | 239 | 0.025576026 | 0.166012853 | 1.67E-05 |
| | Cerebellum | 240 | 0.093581625 | 0.212110632 | 0.001662393 |
| | Cerebellum | 241 | 0.106996004 | 0.273658258 | 0.005058712 |
| | Cerebellum | 243 | 0.040104712 | 0.151942451 | 0.001598643 |
| | Cerebellum | 248 | 0.10391465 | 0.241212581 | 0.003027496 |
| | Cerebellum | 249 | 0.068352422 | 0.212039562 | 0.00177103 |
| | Cerebellum | 250 | 0.020488189 | 0.217056111 | 8.61E-07 |
| | Cerebellum | 256 | 0.053953807 | 0.143541337 | 0.003604706 |
| | medial Frontal gyrus (MFG) | 14 | 0.165683261 | 0.332680307 | 0.003518403 |
| | left dorsal medial Prefrontal cortex (dmPFC) 1 | 145 | 0.119074572 | 0.287968377 | 0.001574639 |
| | left dorsal medial Prefrontal cortex (dmPFC) 2 | 148 | 0.124787167 | 0.264546108 | 0.003726805 |

**Table. 1.** ISC results. All significant nodes, FDR corrected, $q < 0.05$. Light red – significantly more correlated in the right-wing group; Light blue – significantly more correlated in the left-wing group; bold nodes – were involved in processing the stimulus in both groups but were significantly more correlated in one of them.
